# Overdispersed gene expression characterizes schizophrenic brains

**DOI:** 10.1101/441527

**Authors:** Guangzao Huang, Daniel Osorio, Jinting Guan, Guoli Ji, James J. Cai

## Abstract

Schizophrenia (SCZ) is a severe, highly heterogeneous psychiatric disorder with varied clinical presentations. The polygenic genetic architecture of SCZ makes identification of causal variants daunting. Gene expression analyses have shown that SCZ may result in part from transcriptional dysregulation of a number of genes. However, most of these studies took the commonly used approach—differential gene expression analysis, assuming people with SCZ are a homogenous group, all with similar expression levels for any given gene. Here we show that the overall gene expression variability in SCZ is higher than that in an unaffected control (CTL) group. Specifically, we applied the test for equality of variances to the normalized expression data generated by the CommonMind Consortium (CMC) and identified 87 genes with significantly higher expression variances in the SCZ group than the CTL group. One of the genes with differential variability, *VEGFA*, encodes a vascular endothelial growth factor, supporting a vascular-ischemic etiology of SCZ. We also applied a Mahalanobis distance-based test for multivariate homogeneity of group dispersions to gene sets and identified 19 functional gene sets with higher expression variability in the SCZ group than the CTL group. Several of these gene sets are involved in brain development (e.g., development of cerebellar cortex, cerebellar Purkinje cell layer and neuromuscular junction), supporting that structural and functional changes in the cortex cause SCZ. Finally, using expression variability QTL (evQTL) analysis, we show that common genetic variants contribute to the increased expression variability in SCZ. Our results reveal that SCZ brains are characterized by overdispersed gene expression, resulting from dysregulated expression of functional gene sets pertaining to brain development, necrotic cell death, folic acid metabolism, and several other biological processes. Using SCZ as a model of complex genetic disorders with a heterogeneous etiology, our study provides a new conceptual framework for variability-centric analyses. Such a framework is likely to be important in the era of personalized medicine. (313 words)

## Introduction

Schizophrenia (SCZ), one of the most severe psychiatric disorders, affects about 1% of the general population (Knapp et al. 2004; Saha et al. 2005; McGrath et al. 2008). The disorder manifests itself in many different forms and includes both positive behaviors (e.g., delusions, hallucinations, and disorganized speech) and negative behaviors (e.g., absence of reaction, loss of interest in everyday activities, and lack of feeling or emotion). SCZ affects patients differently—people with the disorder vary widely in their symptoms, course of illness and treatment response (Picardi et al. 2012). A reliable clinical typology of SCZ has proved difficult to develop (Kay and Sevy 1990; Andreasen et al. 1997; Budde et al. 2018). Assessment of clinical outcomes in SCZ is challenging (Andreasen et al. 2005; McGrath 2008). Patients diagnosed with SCZ can be classified into those with and without neurodevelopmental impairment (Murray et al. 1992; Kaymaz and van Os 2009; Demjaha et al. 2012). The former category is likely to be due to the impact of risk alleles, copy number variants (CNVs), or early environmental insults such as hypoxic damage to the hippocampus. The latter is more likely to be due to affective dysregulation. Detailed neuroimaging and many measures of psychological variables can also be used to classify SCZ patients into distinct subgroups (Karlsgodt et al. 2010; MacCabe et al. 2012; Arnedo et al. 2015; Cernis et al. 2015). The existence of multiple ways to classify SCZ underscores the marked between-patient variability and within-category heterogeneity associated with the disorder. Thus, SCZ is a highly heterogeneous group of disorders rather than a single disease (Lasalvia et al. 2015; Sommer and Carpenter 2016; van Os 2016).

Although SCZ was described more than 100 years ago, the exact etiology and genetic mechanism of SCZ are still unclear. With an upper bound estimate of heritability of 80% (Hilker et al. 2018), the risk of SCZ is clearly under the substantial genetic influence. Numerous common single nucleotide polymorphisms (SNPs) (Schizophrenia Psychiatric Genome-Wide Association Study 2011; Ripke et al. 2013; Schizophrenia Working Group of the Psychiatric Genomics 2014) and CNVs (International Schizophrenia 2008; Stefansson et al. 2008; Walsh et al. 2008; Xu et al. 2008) have been identified to be associated with the SCZ risk. Nevertheless, like many other complex diseases, SCZ has a polygenic architecture (Schizophrenia Psychiatric Genome-Wide Association Study 2011; Sullivan et al. 2012; Schizophrenia Working Group of the Psychiatric Genomics 2014) and is influenced by environmental factors (Karlsgodt et al. 2010), making it difficult to pinpoint causal mutations.

Gene expression data of SCZ patients has been collected and integrated into analyses (Fillman et al. 2013; Sanders et al. 2013; Fromer et al. 2016; Gusev et al. 2018; Pardinas et al. 2018), in order to improve mechanistic interpretations of risk alleles or directly identify dysregulated genes in relevant tissues. Most studies of SCZ transcriptome adopt the method of differential expression (DE) aimed at the identification of genes expressed significantly higher or lower in SCZ patients than unaffected controls (CTL). However, given the fact that SCZ is highly heterogeneous, we argue that it may not be sufficient to treat SCZ patients as a homogenous group of individuals and expect gene expression in all SCZ samples, compared to CTL samples, is consistently up- or down-regulated. Indeed, a robust overlap between sets of DE genes identified in different SCZ transcriptome studies has not been observed. Many of the statistically significant DE genes cannot be individually connected with any of the current pathophysiological hypotheses of the disease either.

Here, we consider a complementary alternative that disease-relevant genes are expressed more variably in SCZ patients than CTL individuals and therefore lead to an ‘overdispersion’ in gene expression in SCZ. The rationale behind our argument is that SCZ patients are a heterogeneous group of individuals—it is not simply that they share few or no symptoms in common (Andreasen 1999); rather, etiologically, every SCZ patient is ‘ill’ in his or her own way. This can be summarized into the ‘Anna Karenina principle’ for SCZ, in which gene expression of affected individuals varies more than that of healthy individuals, matching with Leo Tolstoy’s dictum that ‘happy families are all alike; each unhappy family is unhappy in its own way.’ We set out to test the hypothesis that overdispersed gene expression in SCZ is a common and important consequence of transcriptional dysregulation. The phenomenon should be more pronounced for genes and pathways that underlie SCZ pathogenesis. The pattern is easily missed or discarded by common workflows, e.g., those implemented in the DE analysis.

The structure of this paper is as follows. First, we represent the results of our differential variability (DV) analysis on single genes, showing an overwhelming pattern of increased variability at the single-gene level associated with SCZ. Second, we develop a multivariate DV analysis and apply it to predefined gene sets. We show that a number of gene sets with SCZ related functions have higher expression variability in SCZ. Third, we examine the contribution of common genetic variants to the expression variability. We compare the relative contributions of these variants in SCZ with that in CTL. To the best of our knowledge, this is the first time this new variability-centric analytical framework has been applied in SCZ. We conclude by providing interpretation of our results in the context of gene discovery and implications in the personalized intervention of SCZ.

## Results

### Overdispersed expression in single genes

We obtained the normalized gene expression data from the CommonMind Consortium (CMC) study (Fromer et al. 2016). The data was generated by using RNA sequencing (RNA-seq) from the dorsolateral prefrontal cortex of people with SCZ and unaffected controls (CTL). We extracted the protein-coding gene expression data for 212 SCZ and 214 CTL samples, all derived from individuals with European ancestry. All analyses of our study were done with this data set. We focused on autosomal protein-coding genes. For each gene, we used the Brown–Forsythe (B–F) test (Brown and Forsythe 1974) to determine whether there is a significant difference in group variances between SCZ and CTL.

We identified 88 of so-called differentially variable (DV) genes at 5% false discovery rate (FDR) level. Among them, 87 show greater expression variances in SCZ than CTL; only one gene (*TAMM41*) shows a smaller variance in SCZ (**Supplementary Table S1**). Thus, at the single-gene level, more dispersed gene expression is an overwhelming feature of SCZ. This pattern is robust against ‘global outliers.’ That is, the pattern is not due to the existence of few outlying individuals with extreme expression values in a large number of genes. To illustrate this, we selected representative DV genes and plotted their expression values across CTL and SCZ samples (**Supplementary Fig. S1**). The expression profiles indicate that there are no SCZ outliers, whose expression is consistently high or low across multiple genes. Also, using one of the DV genes, *ZBTB24*, as an example, we intentionally removed two SCZ samples with the highest and lowest expression to assess the impact of the two outliers on the significance of statistical test for the DV gene detection. The removal of these two extreme samples has almost no influence on the significance level of B–F test (**Supplementary Fig. S2**). These results suggest that increased gene expression variability in SCZ is not driven by a small number of SCZ samples with extremely high or low expression levels. Instead the pattern is due to a systematic overdispersion of gene expression among all SCZ samples.

SCZ DV genes are involved in a variety of biochemical pathways and diverse cellular functions. For instance, *BDNF* is a member of the neurotrophin family of growth factors related to the canonical nerve growth factor; *NECTIN2* encodes a single-pass type I membrane glycoprotein implicated in Alzheimer’s disease; *AIF1* encodes allograft inflammatory factor found in activated macrophages in tissues with inflammation. Furthermore, genetic variants located in or near *HPS5*, *STAR*, *TMEM125*, *RPN2*, and *DNAH1* have been found to be associated with SCZ (Wang et al. 2010; Shi et al. 2011; Goes et al. 2015; Autism Spectrum Disorders Working Group of The Psychiatric Genomics 2017; Lencer et al. 2017).

To validate our results, we used an independent expression data set, which was synthesized from several microarray data sets across multiple studies of neuropsychiatric disorders (Gandal et al. 2018). We applied the same detecting procedure (i.e., B–F test with 5% FDR) to this data set, and identified 11 genes with higher expression variance in SCZ than CTL, with no genes showing the opposite pattern (**Supplementary Table S2**). *VEGFA* is the only gene that was identified using both the CMC and microarray data sets (**Fig. 1A, B**). The gene encodes vascular endothelial growth factor (VEGF), which is a signal protein that stimulates vasculogenesis and angiogenesis (Misiak et al. 2018), playing an important role in neurogenesis, neuronal differentiation, and neuroprotection and regeneration of central nervous system (CNS) cells (Jin et al. 2002; Sun et al. 2003; Howell and Armstrong 2017).

**Fig. 1.**
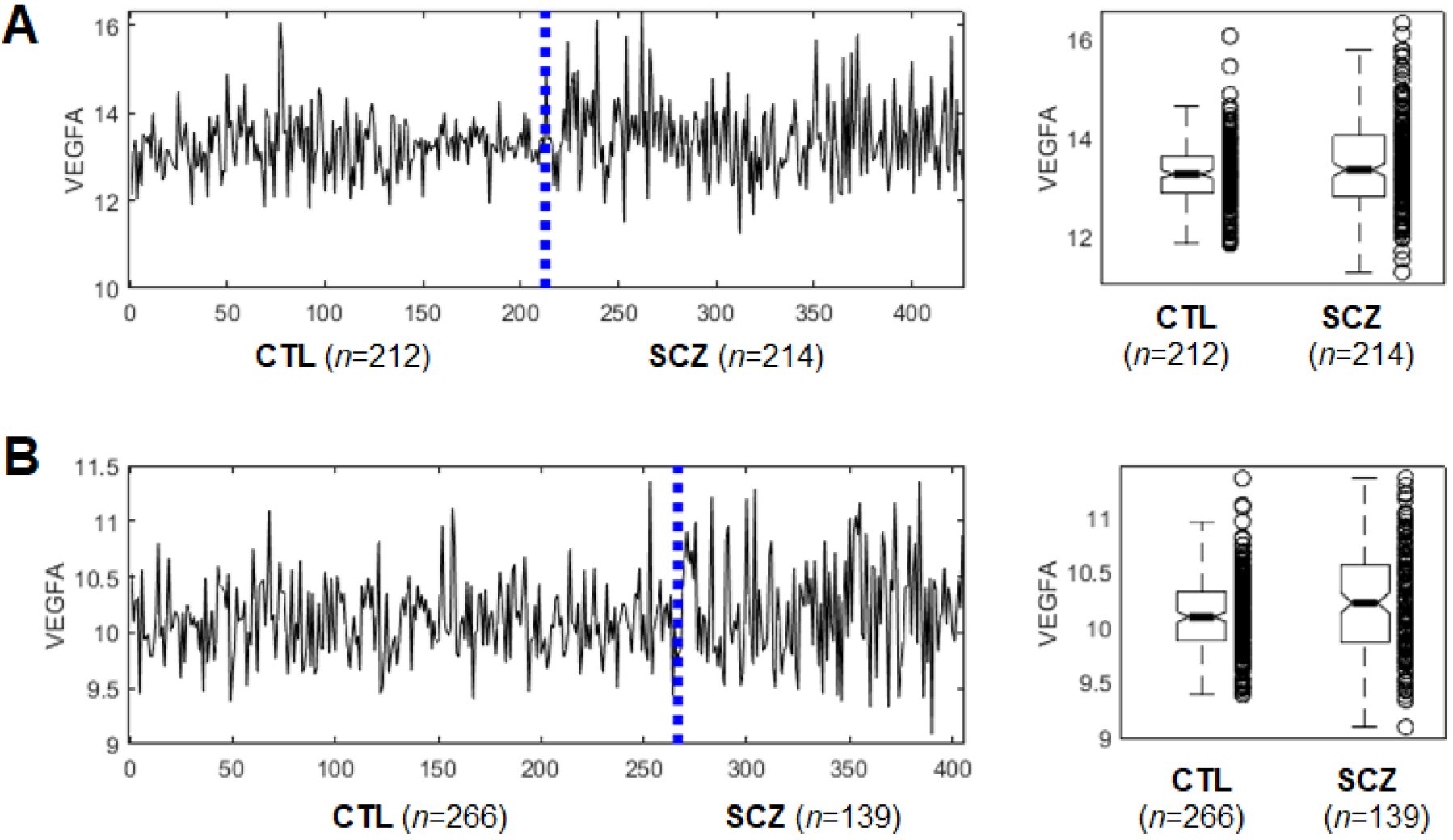
Expression profiles of *VEGFA* show more dispersed expression in SCZ than CTL. (**A**) Normalized expression values in 212 CTL and 214 SCZ samples generated by the CMC study. B–F test *p*-value = 3.6e-6. (**B**) Normalized expression values in 266 CTL and 139 SCZ samples as reported in (Gandal et al. 2018). B–F test p-value = 1.7e-9.

### Overdispersed expression in gene sets

Next, we extend our test for difference in gene expression variances to higher dimensions. To do so, we turned our focus from single genes to gene sets. We adapted the procedure, proposed by Anderson (2006), for analyzing multivariate homogeneity of group dispersions. The procedure is a multivariate analog of Levene’s test for homogeneity of variances and has been widely used in ecology, e.g., in defining the variability in species composition. The modification we made was to use Mahalanobis distance (MD) to replace Euclidian distance originally used in the procedure (see **Methods** for details). This modification is essential to account for collinearity in gene set expression (Zeng et al. 2015; Brinkmeyer-Langford et al. 2016). We applied our MD-based, extended procedure to gene sets, derived from GO ontologies of biological process and molecular function. We identified 19 gene sets with greater multivariate expression variance in SCZ, all with a nominal *p*-value < 0.05 (**Table 1**). Among these gene sets, four are related to brain development: (1) *cerebellar cortex formation*, (2) *cerebellar cortex morphogenesis*, (3) *cerebellar Purkinje cell layer development*, and (4) *neuromuscular junction development*. The first three sets share 12 common genes: *AGTPBP1*, *ATP2B2*, *ATP7A*, *CACNA1A*, *CEND1*, *DLL1*, *FAIM2*, *HERC1*, *LDB1*, *LHX1*, *LHX5* and *RORA;* the last gene set shares no gene with any of the first three. These results suggest that SCZ brains are characterized by abnormally high expression variance in brain development genes. This finding is consistent with the general consensus of that SCZ is a brain disorder with structural and functional changes in the cortex and the connections between different cortical regions (Karlsgodt et al.2010; Demjaha et al. 2012; Kelly et al. 2018). In addition to these four gene sets pertaining to brain development, several other gene sets unrelated to the CNS, but likely to be implicated in SCZ are highlighted here. These include *necrotic cell death* (Jarskog et al. 2005; Catts and Weickert 2012), *regulation of endoplasmic reticulum unfolded protein response* (Wang and Kaufman 2012), and *folic acid-containing compound metabolic process* (Saedisomeolia et al. 2011; Roffman et al. 2013; Brown and Roffman 2014). For each significant gene set, we use the method of Garthwaite–Koch partition to estimate the relative contribution of each gene in the gene set to MD (Garthwaite and Koch 2016). We present top five genes that contribute most to MD for each gene sets in **Table 1**.

**Table 1.**
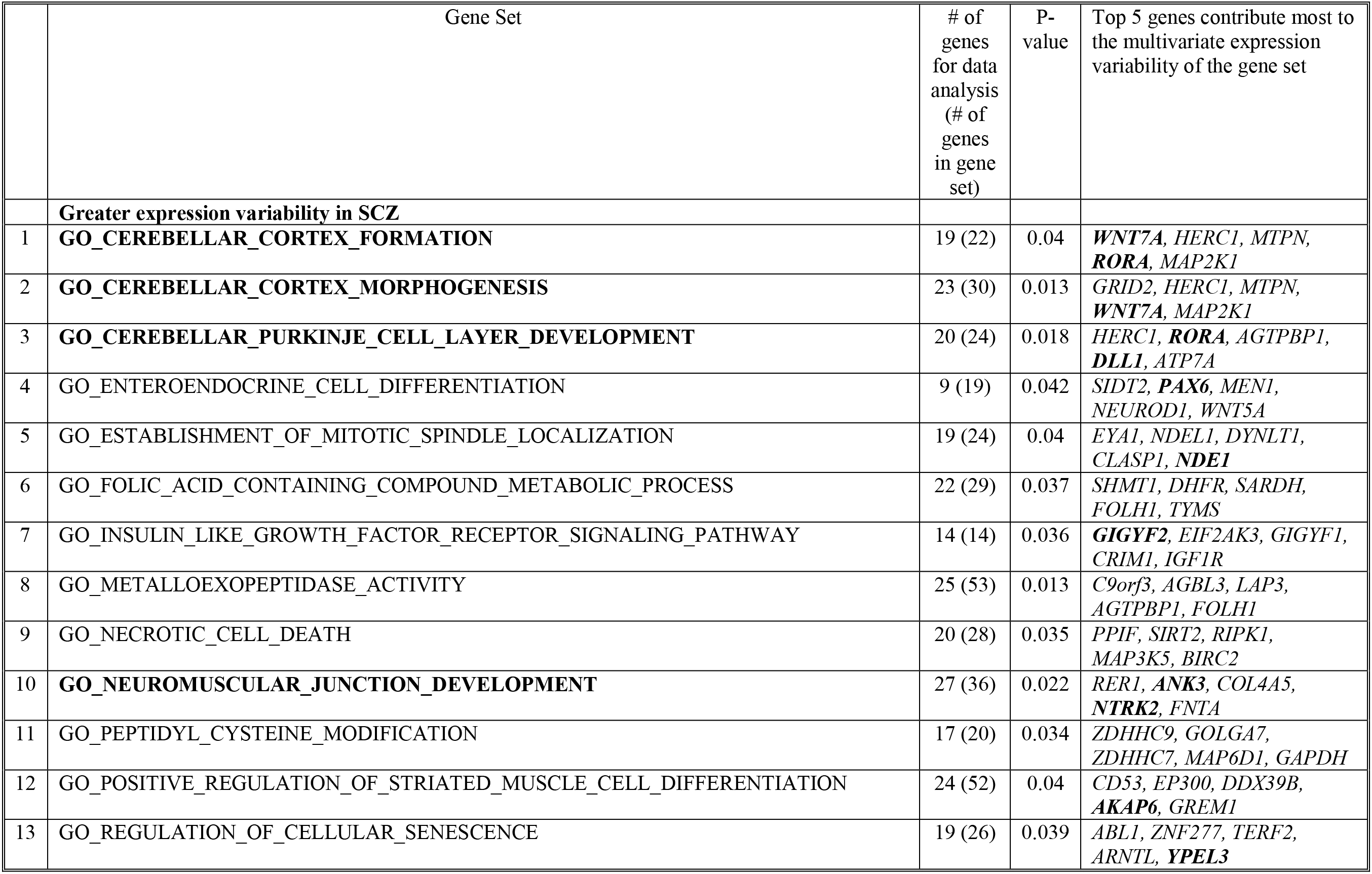
Functional gene sets showing differential expression variability between SCZ and CTL. For each gene sets, top five genes that contribute most to expression variability are given. Genes highlighted with bold font are those known to be implicated in SCZ.

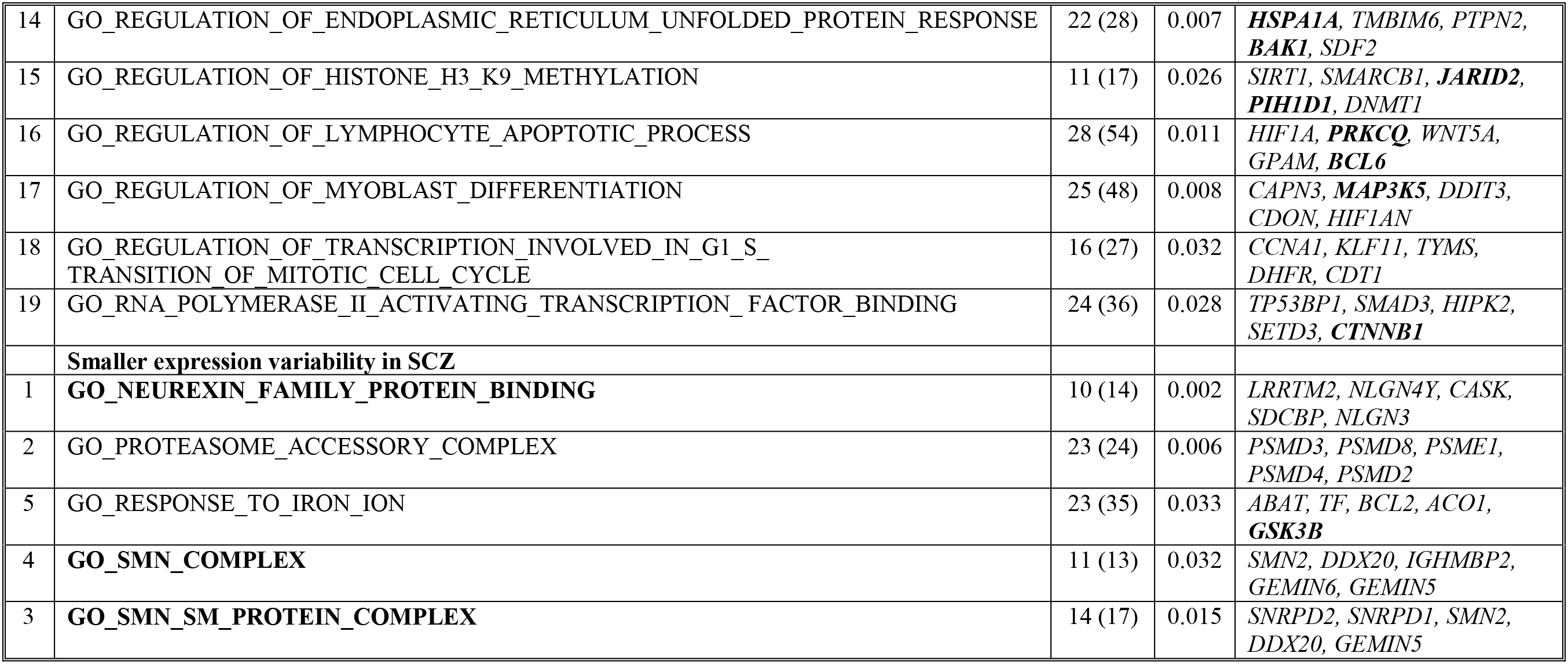

To illustration the overdispersion pattern of gene expression in SCZ, we used the gene set of *cerebellar cortex morphogenesis* as an example (**Fig. 2**). We extracted the expression data of 23 genes in the gene set, pooled SCZ with CTL samples, and performed principal component (PC) analysis. On the first and second PC space, SCZ samples are more dispersedly distributed than CTL samples (**Fig. 2A, B and C**). Accordingly, the variance in MD of individual SCZ samples to the centroid is significantly greater than that of CTL samples (**Fig. 2D**).

**Fig. 2.**
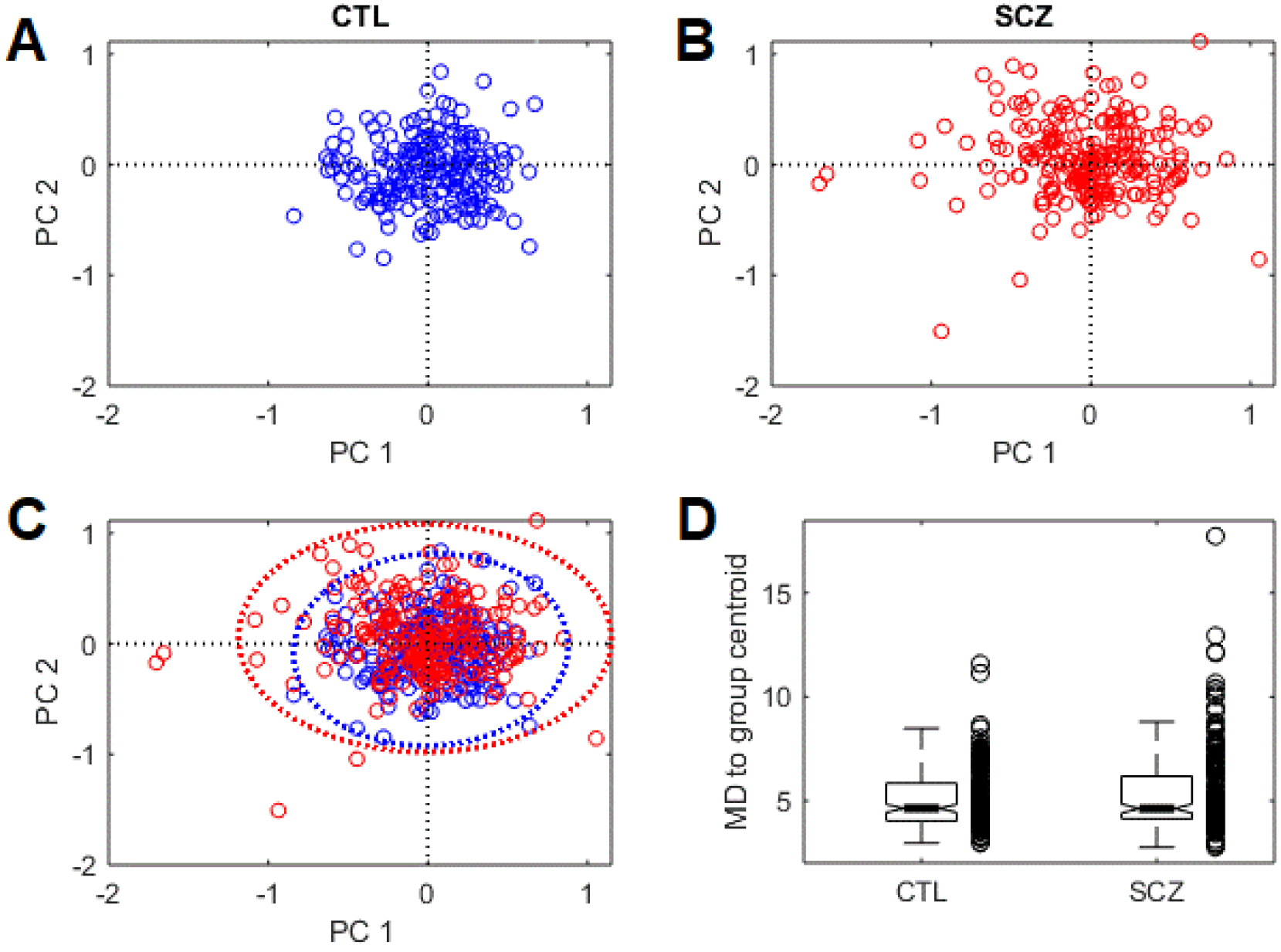
Gene set, *cerebellar cortex morphogenesis*, show more dispersed expression in SCZ. The PCA analysis was performed with the gene set expression matrix of pooled samples that contain all SCZ and CTL samples. (**A**) Distribution of CTL samples on the PCA space defined by the first two PCs. SCZ samples are made invisible by plotting in white color; (**B**) Distribution of SCZ samples with CTL samples made invisible. (**C**) Distribution of all samples in the PCA space. Dashed lines indicate the 99% confidence ellipses. (**D**) Boxplot of MD vectors in SCZ and CTL groups, showing the high within-group variance in SCZ.

We also identified five gene sets showing the opposite pattern, i.e., smaller expression variance in SCZ than CTL. Among these five, two gene sets contain largely overlapped genes encoding proteins of the survival motor neuron (SMN) complex, which plays a role in neuronal migration and differentiation (Giavazzi et al. 2006), with a potential role in SCZ (Comley et al. 2016; McLaughlin et al. 2017). One gene set consists of genes that encode synaptic cell adhesion proteins, interacting with neurexins, which is essential for brain function (Missler et al. 2012).

### Statistical power analysis

We used computer simulations to conduct a power analysis for our extension of Anderson’s procedure. The simulations were done with combinations of a series of sample size and varying levels of variance difference between case and control groups (see **Methods** for details). We considered a balanced design with the sample size of case group equals to that of the control group. The result of the power analysis shows that the power of our extended test starts to increase when the sample size per group is over 800 (**Supplementary Fig. S3**). The test becomes highly sensitive when the sample size per group reaches 1000—in this case, when variances of two groups differ by two-fold, the statistical power of our test can reach 80%. The power analysis suggests that the sample size we used in real data analysis (212 and 214 for SCZ and CTL, respectively) is too small to produce any meaningful results, based on the data generation algorithm we used for the simulations. On the other hand, our power analysis suggests that the results we obtained in real data analysis (**Table 1**), albeit none of the gene sets survived the multiple test correction, are unexpected by chance. We are inclined to think that it was that the overdispersion pattern in multivariate gene expression in SCZ itself is strong enough to be captured by our test even with limited numbers of samples.

### Common genetic variants contribute to dispersed gene expression in SCZ

To assess the contribution of genetic variants to the gene expression dispersion in SCZ, we conducted an expression variability QTL (evQTL) mapping analysis. An evQTL is a SNP polymorphism whose genotypes are associated with different group variance in gene expression (Hulse and Cai 2013; Wang et al. 2014b). In this case-control study setting, we were more interested in SNPs with different effects on SCZ and CTL, and thus we set out to identify SCZ-specific evQTLs. We tested all common SNPs segregating in SCZ and CTL, i.e., minor allele frequency (MAF) > 0.15 in both populations, to identify those with genotypes associated with gene expression variance in SCZ (*p* < 1e-7, B–F test) but not in CTL (*p* > 0.05, B–F test, **Fig. 3A**). We identified 2,503 SCZ-specific evQTLs involving 1,453 distinct autosomal protein-coding genes (**Supplementary Table S3**, see **Supplementary Fig. S3** for more examples). For comparison, we used the same procedure and *p*-value cutoffs to identify CTL-specific evQTLs with SCZ samples as the background group. We identified 2,076 CTL-specific evQTLs involving 1,277 genes. Although the number of CTL-specific evQTLs is comparable to that of SCZ-specific evQTLs, a q-q plot shows that the overall statistical significance is much stronger for SCZ-specific evQTLs, especially for those highly significant ones (**Fig. 3B**). We plotted the links between these highly significant evQTLs (*p*<1e-9) with their target genes for both SCZ and CTL results. Most of these relationships are *trans*-acting and more SCZ-specific evQTLs connect with target genes and form a denser picture than CTL-specific ones do (Fig. 3C).

**Fig. 3.**
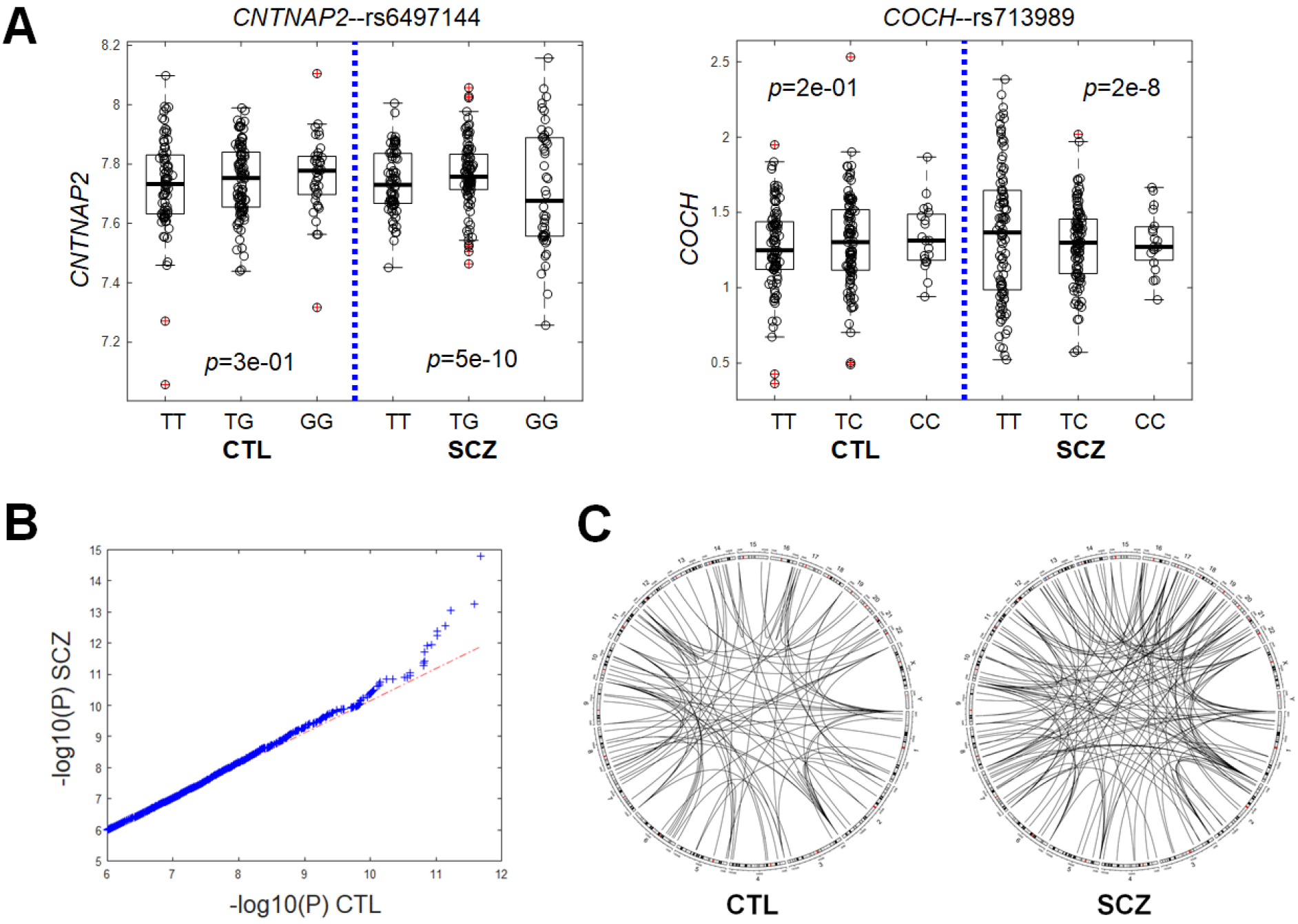
Genetic variants associated with gene expression variability in SCZ. (**A**) Two examples of SCZ-specific evQTLs, showing significant differences in expression variances between genotypes in SCZ but not in CTL. P-values of B–F test for three genotype groups in SCZ and CTL, respectively, are given. (**B**) Quantile-quantile plot of *p*-values for evQTL associations in SCZ (y-axis) against those in CTL (x-axis). (C) Difference in density of highly significant evQTL associations between variants and genes in SCZ and CTL.

## Discussion

Our analysis focuses on the differences in expression variances between groups. The DV approach we adopted here could be rooted in the theory of dynamics of correlation and variance in systems under the load of environmental factors (Gorban et al. 2009). For heterogonous disorders like SCZ, we show that the DV approach is powerful in identifying significant genes and characterizing biological pathways and processes critical to the disorder. DV approach is complementary to the more commonly used DE method. So far, identified DE genes between SCZ and CTL show a tremendous functional diversity and often fail to form supported, functionally interpretable gene sets or pathways, which is not unexpected under the assumption of substantial heterogeneity in SCZ pathophysiology (Sanders et al. 2013).

Overdispersion is the term we borrow from statistics to describe the presence of greater variability (statistical dispersion) in gene expression data in SCZ than would be expected based on that in CTL. The overdispersion pattern we observed in SCZ is in line with the fact that clinical heterogeneity amongst people diagnosed with SCZ is high, which has hampered the development of individual treatment and research into new treatment strategies. We assume that key pathways underlying SCZ risk are disrupted via many different causes—genetic, epigenetic, and environmental. The consequence of these disruptions is collectively reflected as dysregulated gene expression, which is in turn characterized by an increased level of group dispersions (variances). Thus, the degree of gene expression dispersion is an excellent predictor of functional disruptions. Even if the illness of SCZ in every affected individual arises from a different specific cause, each will nonetheless share disruption of related key biological processes. Through assessing the expression dispersion in genes, we identify the related pathways dysregulated in patients affected by SCZ.

In this study, *VEGFA*, the gene encoding vascular endothelial growth factor, stands out as the most significant single DV gene, cross-validated with two independent expression data sets. The discovery of this significant gene is consistent with accumulating evidence indicating that SCZ is accompanied by abnormal vascularization (Hanson and Gottesman 2005; Moises et al. 2015; Misiak et al. 2018). The role of VEGF in causing neurovascular dysfunction has been known to be correlated with hypoxia-ischemia insults during early life and is strongly associated with cognitive dysfunction (Howell and Armstrong 2017; Misiak et al. 2018). Clinical studies examining peripheral VEGF levels in SCZ versus CTL have yielded conflicting results. While some studies found elevated serum VEGF concentrations in individuals with SCZ (Pillai et al. 2016; Balotsev et al. 2017), others revealed no significant difference (Di Nicola et al. 2013; Murphy et al. 2014; Haring et al. 2015; Lizano et al. 2016), or lower concentrations (Lee et al. 2015; Xiao et al. 2018). The extremely high between-individual variability in *VEGFA* expression may explain such conflicting findings, although the effect of antipsychotic agents on plasma VEGF level cannot be ruled out [(Pillai and Mahadik 2006; Lee et al. 2015) *cf*. (Haring et al. 2015)].

One of the contributions of this study is to develop the homogeneity test of multivariate dispersions further using an MD-based extension. The initially proposed procedure by Anderson (2006) is flexible enough to allow any distance measure to be adapted. We have previously used a slightly different implementation of the test to identify dysregulated gene sets in autism spectrum disorder (Guan et al. 2016; Guan et al. 2018). Other implementations include the one based on the means of within-group distances, which does not require group center calculations to obtain the distance statistic (Gijbels and Omelka 2013) and the one to compare *k* populations based on Fréchet variance for general metric space valued data objects, with emphasis on comparing means and variances (Dubey and Müller 2017). The most important feature shared by these tests is their capability of capturing the between-group difference in multivariate covariance of variables. Our method, when applied to gene sets, has approved to be useful in identifying functionally meaningful gene sets. Our results reveal that transcriptional dysregulation in genes responsible for brain development is significantly implicated in SCZ, which is in line with the consensus of SCZ being a brain disorder (Karlsgodt et al. 2010; Demjaha et al. 2012; Kelly et al. 2018). Our analysis with gene sets also supplies new insights into other mechanisms that may not have been implicated in SCZ. These include cell death, folic acid metabolism, and metalloexopeptidase activity. We also report the opposite pattern detected in several gene sets, for which expression variability is reduced rather than elevated in SCZ.

In the final section of our analysis, we explore that relationship between genetic variants and expression variability. We are among the earliest adopting the ‘vQTL’ strategy (Ronnegard and Valdar 2011) and have applied it to gene expression data (Hulse and Cai 2013; Brown et al. 2014). In humans, we have identified a significant number of evQTLs (Wang et al. 2014a; Yang et al. 2016). In this study, we emphasize the difference in the statistical significance of evQTLs in SCZ and CTL. We show that evQTLs with higher statistical significance are preferably present in SCZ, suggesting that genetic variants may play a more important role in shaping the gene expression variability in SCZ than in CTL. It is not clear though whether these common genetic variants exert destabilizing function on their own or act through interacting with each other (Yang et al. 2016). Nevertheless, these results have clinical implications. For example, *CALM1*-rs2123259 (GT) and *HTR1A*-rs12440923 (TC) are two identified SCZ-specific evQTLs (**Supplementary Table S3**). *CALM1* encodes for calmodulin 1 and *HTR1A*, serotonin 1A receptor (or 5-HT_1A_ receptor). Both gene products are targets of antipsychotic drugs. For example, aripiprazole is a partial agonist at the 5-HT_1A_ receptor; chlorpromazine binds to calmodulin to exert an inhibitory effect (Marshak et al. 1985) and exhibited antagonist activity at serotonin 1A receptors (Newman-Tancredi et al. 1998). According to the evQTL pattern, the expression level of *CALM1* in SCZ patients with TT genotype at rs2123259 (chr15:87257140_hg19) is highly variable from patient to patient, and the same as for the *HTR1A* expression in patients with CC genotype at rs12440923 (chr15:93164993_hg19). This information ought to be taken into account in antipsychotic medication administration.

Several caveats exist in our study. We could not recover most of our results using the independent expression data set, which is based on microarray technology. This discrepancy might be attributed to the lack of dynamic range of the microarray data compared to the data generated by using RNA-seq. We also note that our DV analysis for both single gene and gene set settings are sensitive to the procedure and method used for processing and normalizing input expression matrix. In this regard, we recognize that the CMC data set has been appropriately processed and carefully normalized, allowing the subtle patterns in residual expression data to be revealed in our analysis. This, of course, sets a high bar for future large-scale SCZ transcriptome projects.

In conclusion, we establish a new analytical method, based on statistical tests for homogeneity in group variances, to reveal overdispersed gene expression in SCZ, which is associated with the heterogeneous nature of this disorder characterized by varying clinic presentations and individualized symptoms. We show our method contributes to the discovery of genetic underpinnings of SCZ that are notoriously difficult to determine. We identify single genes as well as gene sets showing greater gene expression variability in SCZ patients but not in unaffected controls. Functional interpretation of these genes points us to dysregulation of brain function pertaining to a number of new mechanisms. Our evQTL analysis reveals the contribution of common genetic variants to expression variability in SCZ. We anticipate our study inspires new conceptual development toward variability-centric analyses in the era of personalized medicine.

## Materials and Methods

### Data sets

We obtained the gene-level expression data generated by the CommonMind Consortium (CMC) study (Fromer et al. 2016). The expression matrix was derived from the raw read count matrix through a series of normalization and adjustment. Briefly, 16,423 genes with at least one CPM (read counts per million total reads) in at least 50% of the individuals were retained and processed. The initial normalization was done using voom (Law et al. 2014). Weighted linear regression was then performed for each gene to control for known covariates. The data was further adjusted for hidden variables detected by surrogate variable analysis [see (Fromer et al. 2016) for details]. We downloaded the processed data matrix, which was the one used for eQTL identification in the original CMC study (Fromer et al. 2016), from the web page at https://www.synapse.org/#!Synapse:syn5609491 with the file name ‘CMC_MSSM-Penn-Pitt_DLPFC_mRNA_IlluminaHiSeq2500_gene-adjustedSVA-dataNormalization-noAncestry-adjustedLogCPM’. From this downloaded data matrix, we extracted data generated from European ancestry individuals, including 212 SCZ and 214 CTL samples. We used this European data set throughout all subsequent analyses. In addition to the CMC data, an independent, microarray-based gene expression data, collected by a study of multiple psychiatric disorders (Gandal et al. 2018), was obtained. From this microarray data set, we extracted expression data for 139 SCZ and 266 CTL samples.

### Test for homogeneity of expression variances of single genes

Brown–Forsythe (B–F) test was applied to each gene to test for the significant difference in expression variance between SCZ and CTL subjects. B–F test is a statistical analysis related to Levene’s test—both are tests for homogeneity of variance. B–F test involves determining an absolute deviation score from group medians, while Levene’s test from group means. After running the B–F test for all individual genes, we used the Benjamini–Hochberg (B–H) procedure (Benjamini and Hochberg 1995) to control the FDR of hypothesis tests.

### Test for multivariate homogeneity of expression variances of gene sets

Anderson (2006) proposed a distance-based test of homogeneity of multivariate dispersions for a one-way ANOVA design (Anderson 2006). To identify gene sets with differential multivariate expression variability between SCZ and CTL, we adopted the ‘Anderson06’ test and applied it to functional gene sets. These functional gene sets are predefined in the MSigDB v5.2 (Liberzon et al. 2015). We selected those defined with GO ontologies of biological process and molecular function. For a given gene set, we measured multivariate expression dispersion (variance) of genes in SCZ and CTL separately. We calculated Mahalanobis distance (MD) of each SCZ and CTL individuals to their group centroids in the multivariate space. Compared to the Euclidean distance used in the original Anderson06 test, MD is a more appropriate distance metric for gene set expression, because expression levels of different genes in a gene set are likely to be correlated (Zeng et al. 2015; Brinkmeyer-Langford et al. 2016). To test if the dispersions (variances) of SCZ and CTL groups are different, MDs of group members to the group centroid were subject to ANOVA (Anderson 2006). To determine which genes in a gene set have the most influence on the MD, we used the method proposed by Garthwaite and Koch (2016).

### Power analysis of MD-based Anderson06 test

We conducted simulations to evaluate the power of the MD-based Anderson06 test. We fixed the size of the input gene set at 23, based on the number of genes in the actual gene set of *GO_CEREBELLAR_CORTEX_MORPHOGENESIS* (**Table 1**). We also fixed the covariance matrices, ∑_0_ and ∑_1_of the input gene set for CTL and SCZ samples, respectively, using the values calculated from the actual expression data of the gene set. We varied values of two parameters: (1) the sample size per group, and (2)*α*, a factor multiplied to ∑_1_ to increase multivariate variability, and assessed the power as a function of these two parameters. We set the sample size of the SCZ group equals to that of the CTL group. We assume that the gene expression levels of the gene set obey a multivariate normal distribution (MVN). The simulations were done as follows:

Step 1: We calculated the mean vector *μ*_0_ and covariance matrix ∑_0_ from the expression values of the real gene set in CTL, and *μ*_1_ and Σ_1_ in SCZ.

Step 2: We generated the simulated expression data of CTL samples from *MVN*(*μ*_0_, ∑_0_). We generated the expression data of SCZ samples from *MVN*(*μ*_1_, (1 + *e*^*l*^)∑_1_), where *l* obeys uniform distribution (0, *α*).

Step 3: We set *α* = 1,1.2, …, 5 and the sample size *n* = 100,200, …, 2000. For each pair of the two parameters, we simulated expression data matrix X_1_ for SCZ. Accordingly, we simulated expression matrix X_0_ for CTL. For samples in the data X_1_, we calculated their MD to the group centroid and obtained the vector *D*_*1*_ for SCZ. In the same way, we obtained the vector *D*_*0*_ for CTL. We used ANOVA to compare the difference between *D*_*1*_ and *D*_*0*_. For each pair of *α* and *n*, this process was repeated for 100 times and the number of ANOVA *p* < 0.01 was recorded as the probability of that the test correctly rejects the null hypothesis.

### Identification of SCZ-specific evQTLs

The genotype data of the SCZ and CTL samples was obtained from the CMC Knowledge Portal web site at https://www.synapse.org//#!Synapse:syn3275211. For each SNP, the MAF is calculated for SCZ and CTL samples separately. The SNPs with both MAF > 0.15 were retained for the analysis. B–F tests were conducted with three genotype groups of each SNP to examine whether there is a significant difference in expression variances between any two groups. The B-F test was conducted for SCZ and CTL samples separately. The SCZ-specific evQTLs were called when the *p*<1e-7 in SCZ samples and *p*>0.05 in CTL samples; the CTL-specific evQTLs were called when the *p*<1e-7 in CTL samples and *p*>0.05 in SCZ samples. SZGR 2.0 (Jia et al. 2017) was used to identify antipsychiatric drugs whose targets are evQTL genes.

### Software

Source code to deterministically generate all results and plots in this paper can be found at https://github.com/cailab-tamu/scz_ge_dispersion.

## Acknowledgments

We thank Michael Gandal for sharing the processed microarray data, Barbara Gastel for critically reading and editing the manuscript, and Ahmad Kawam for helping with methodology development. We acknowledge the Texas A&M Institute for Genome Sciences and Society (TIGSS) for providing computational resources. JJC was partially supported by TAMU T3 seed grant and NIH grant R21AI126219. This work was also supported by the National Natural Science Foundation of China (No. 61573296 and 61803320) and the fund of China Scholarship Council to GH. The main data set used for the analyses in this study was generated as part of the CommonMind Consortium supported by funding from Takeda Pharmaceuticals Company Limited, F Hoffman-La Roche and NIH Grants R01MH085542, R01MH093725, P50MH066392, P50MH080405, R01MH097276, RO1-MH-075916, P50M096891, P50MH084053S1, R37MH057881 and R37MH057881S1, HHSN271201300031C, AG02219, AG05138 and MH06692. Brain tissue for the study was obtained from the following brain-bank collections: the Mount Sinai NIH Brain and Tissue Repository, the University of Pennsylvania Alzheimer’s Disease Core Center, the University of Pittsburgh NeuroBioBank and Brain and Tissue Repositories and the NIMH Human Brain Collection Core. CMC Leadership: P Sklar, J Buxbaum (Icahn School of Medicine at Mount Sinai), B Devlin, D Lewis (University of Pittsburgh), R Gur, C-G Hahn (University of Pennsylvania), K Hirai, H Toyoshiba (Takeda Pharmaceuticals Company), E Domenici, L Essioux (F Hoffman-La Roche), L Mangravite, M Peters (Sage Bionetworks), T Lehner, B Lipska (NIMH).

## Supplementary Figure Legends

**Supplementary Fig. S1.**
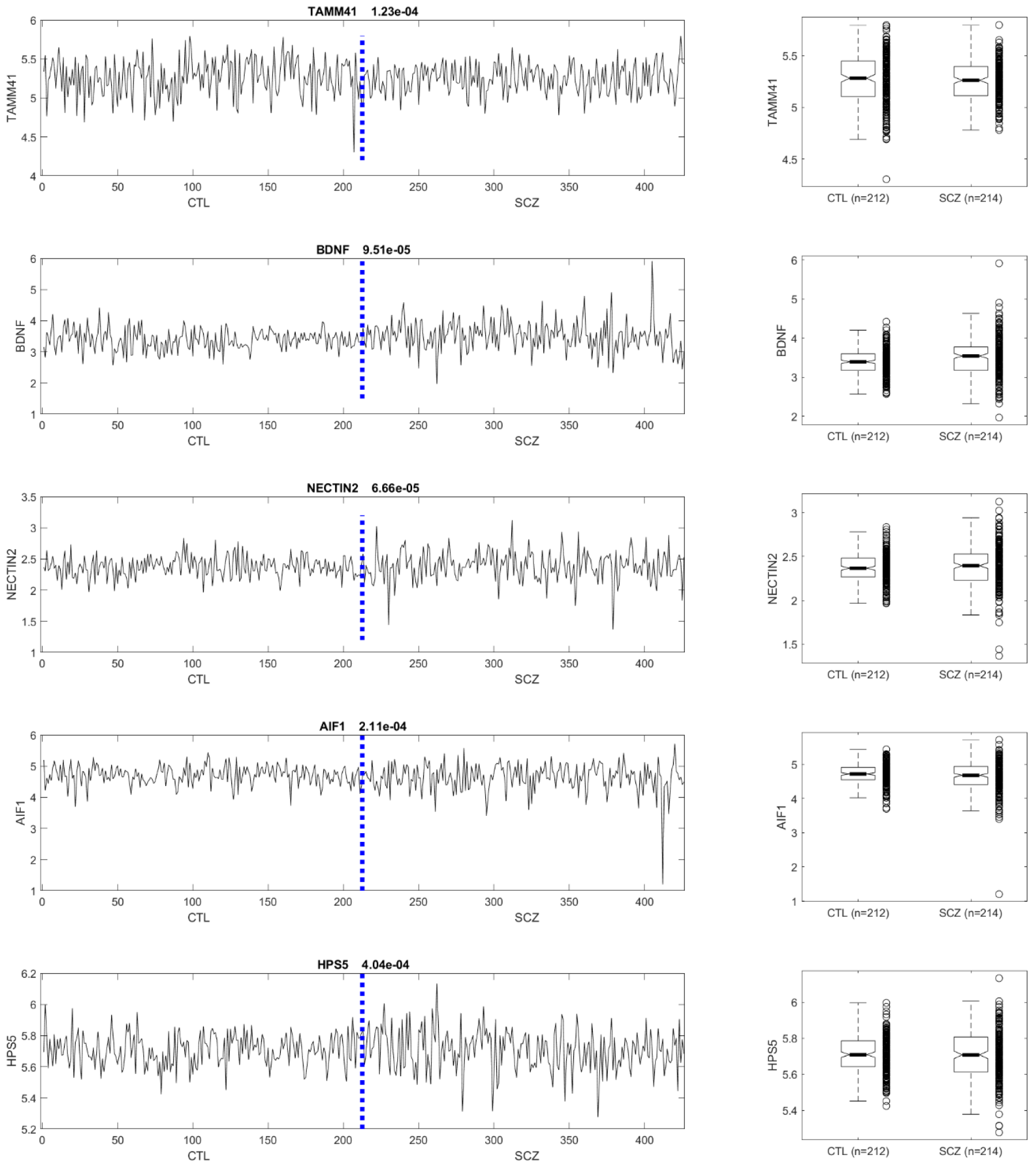
Expression profiles for selected genes with expression variance in SCZ is significantly different from that in CTL (FDR<0.05).

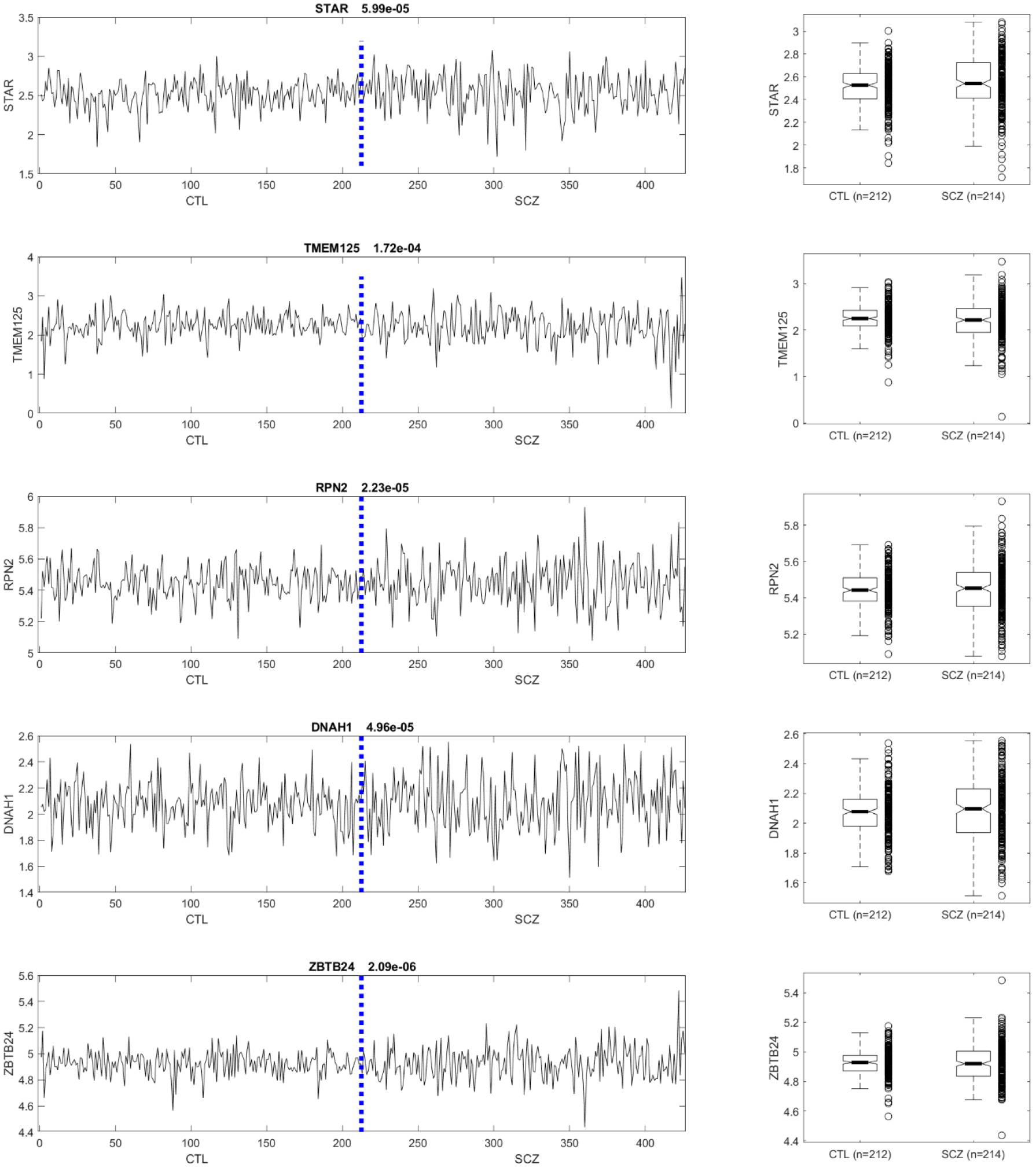

**Supplementary Fig. S2.**
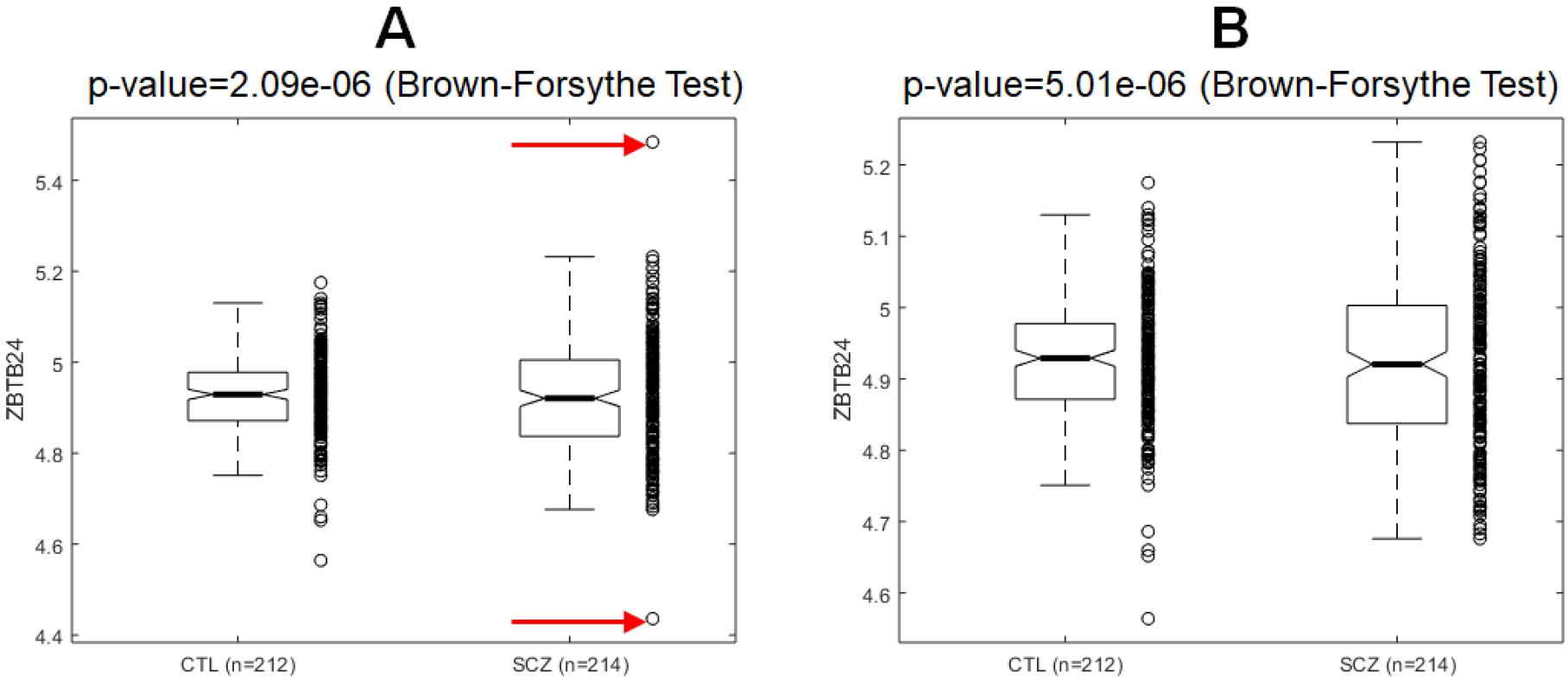
Removing ‘outlier’ samples in the SCZ group does not cause substantial change in the significance level of B–F test. (**A**) Red arrows indicate the two outliers in the original SCZ group that are removed in the secondary analysis. (**B**). The result of secondary analysis showing the significance remains after the two outliers are removed. The overall variance in the SCZ group is significantly greater than that in the CTL group.

**Supplementary Fig. S3.**
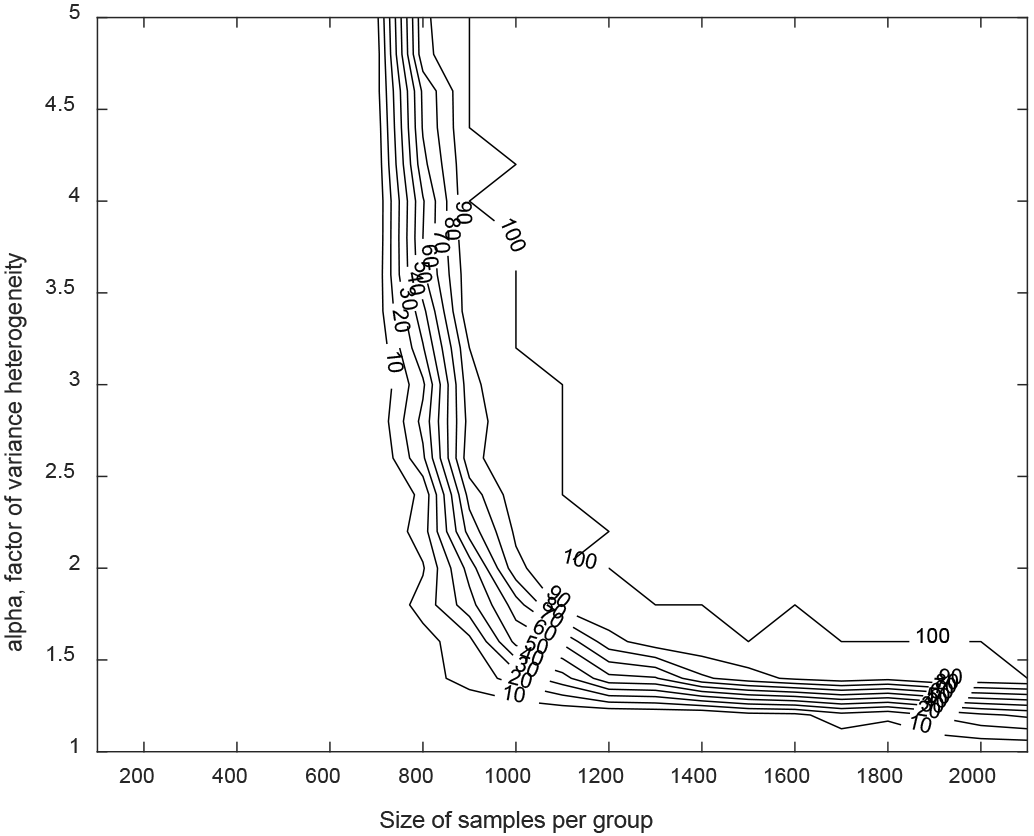
Statistical power of the MD-based Anderson (2006) test to detect significant heterogeneity in multivariate gene expression between two groups. Each contour line depicts the power, i.e., the likelihood level (%) that a random data set would yield a chance finding, as a function of the sample size per group, and alpha—a factor for introducing variance heterogeneity to one of the groups.

**Supplementary Fig. S4.**
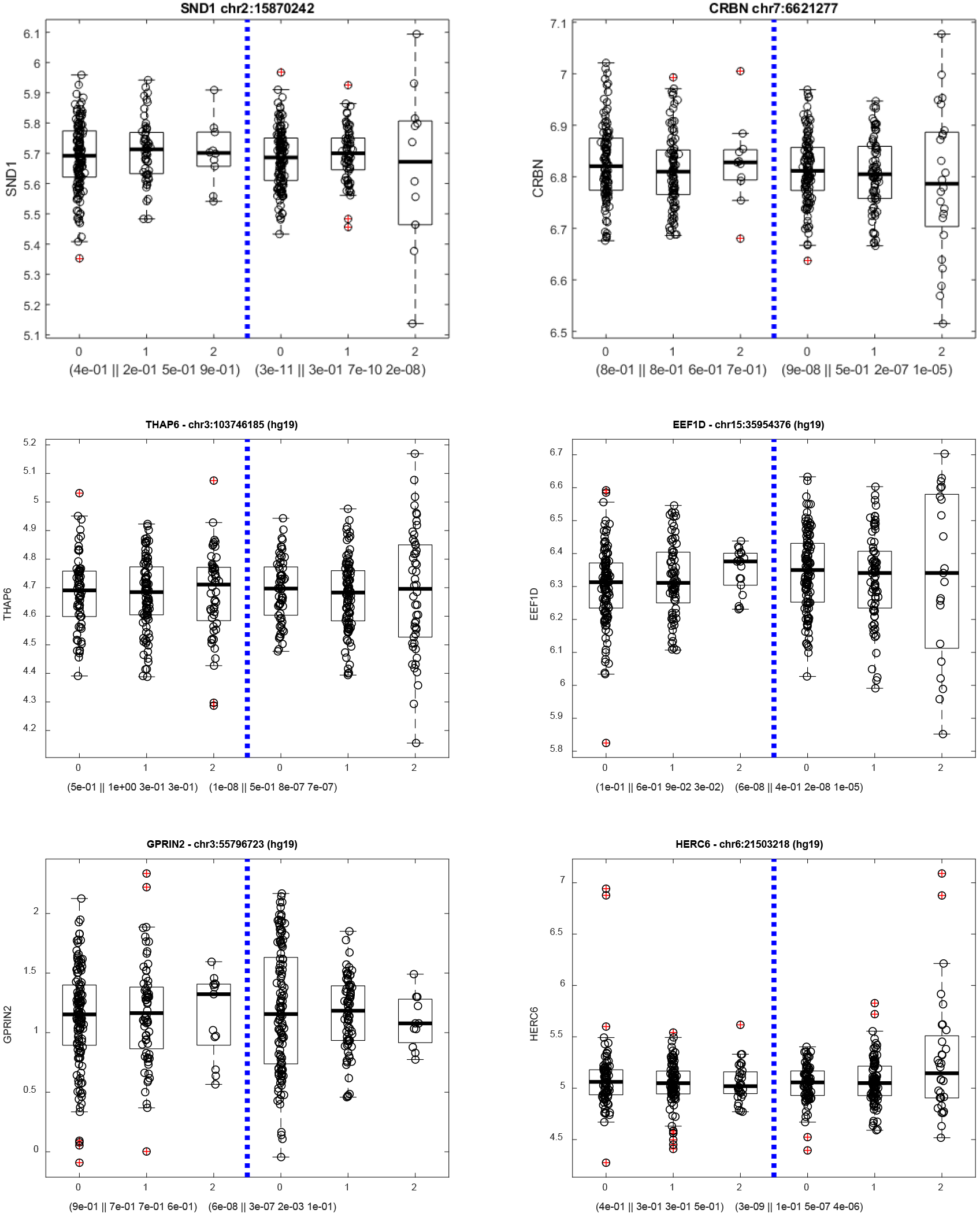
Additional SCZ-specific evQTL examples.

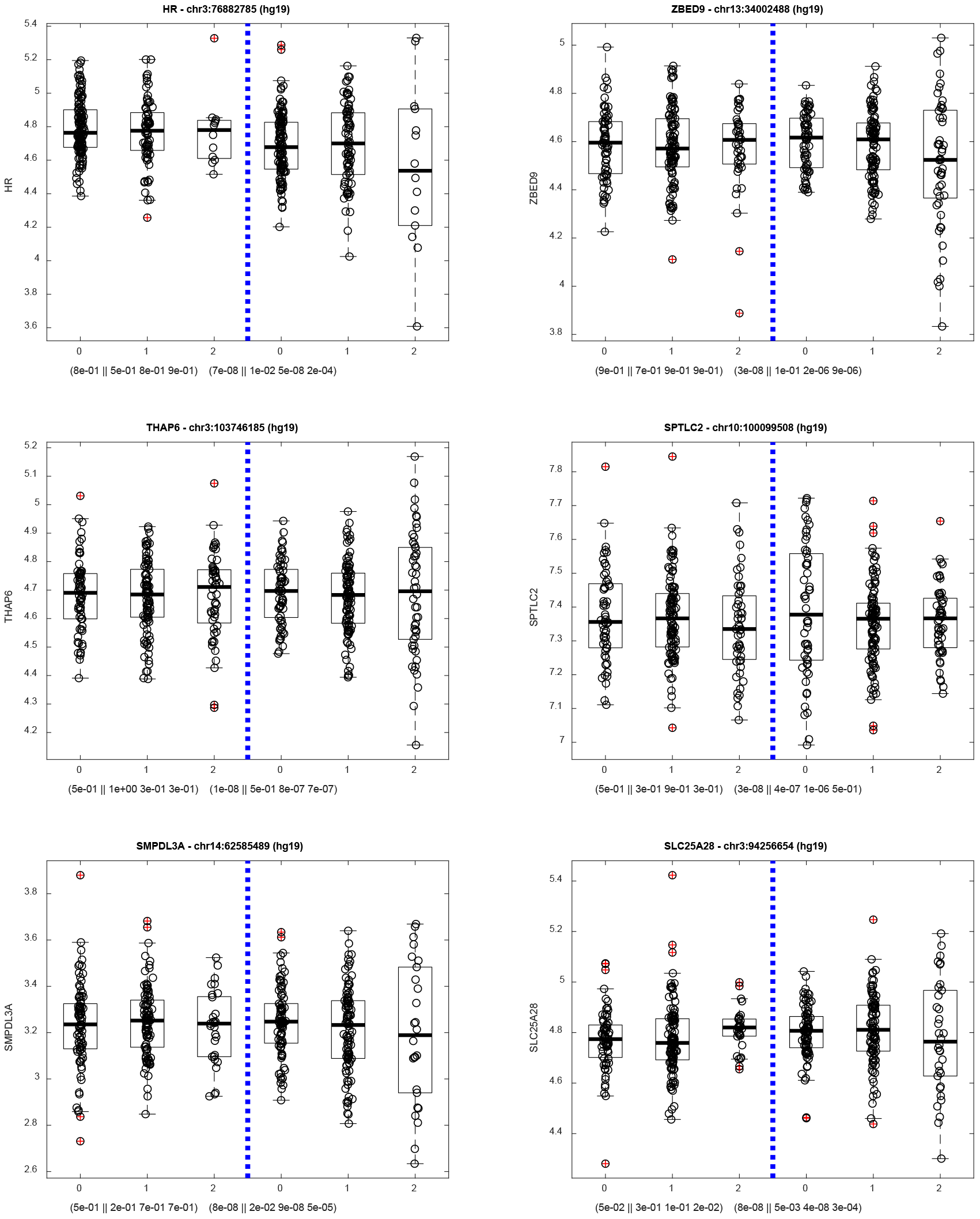

### Supplementary Table Legends

**Supplementary Table S1.** List of 88 differentially variable (DV) genes whose expression variance in SCZ is significantly different from that in CTL (FDR<0.05). For each gene, the P-value of Brown–Forsythe (B–F) test, comparison between expression STD in SCZ and CTL, and gene name, are given.

**Supplementary Table S2.** List of 11 differentially variable (DV) genes identified using microarray gene expression data set of the study of (Gandal, Haney et al. 2018). For each gene, gene symbol, P-value of Brown–Forsythe (B–F) test, and comparison between expression STD in SCZ and CTL, are given.

**Supplementary Table S3.** List of 2,530 SCZ-specific evQTLs. Name of the target gene, genomic position (hg19) of SNP, p-values of B-F test in CTL and SCZ, are given.

